# Is intron removal by recursive splicing necessitated by the evolution of genomic gigantism?

**DOI:** 10.1101/2024.12.11.627982

**Authors:** Alexander Nichols Adams, Rachel Lockridge Mueller

## Abstract

Splicing of introns is an important step that pre-mRNA transcripts undergo during processing in the nucleus to become mature mRNAs. Although long thought to occur in a single step, introns are now also known to be removed in multiple steps through a process called recursive splicing. This non-canonical form of splicing is hypothesized to increase splicing fidelity, particularly in longer introns. In species that have evolved gigantic genome sizes, overall intron lengths are much longer than in genomes of typical size. Using West African lungfish (*Protopterus annectens*; genome size ∼40Gb) as a model, we use total RNA-seq data to test the hypothesis that gigantic genomes have shifted to rely predominantly on recursive splicing to manage their long introns. Our results suggest levels of recursive splicing at conserved sites in the lungfish that are similar to those seen in humans, indicating that genome-wide intronic expansion accompanying genomic gigantism does not necessitate ubiquitous use of recursive splicing. However, in addition to these results, we also observed patterns of decreasing RNA-seq read depths across entire intron lengths and note that both canonical co-transcriptional splicing and stochastic recursive splicing using many random splice sites could produce this pattern. Thus, we infer canonical co-transcriptional splicing and/or stochastic recursive splicing – but not ubiquitous recursive splicing at conserved sites – manage the removal of long introns in gigantic genomes. We conclude that the vast increase in intron length in gigantic genomes does not appear to have coincided with a fundamental shift in splicing mechanisms.

**Statement and Declarations:** The authors declare no competing interests.

## Introduction

As genomes evolve towards larger sizes, the amount of protein-coding DNA does not increase in proportion to overall genome size (Hedges and Kumar 2002; Smith et al. 2009; Wang et al. 2021). Instead, genomic expansion reflects a relative increase in transposable elements and intronic sequence (Sun et al. 2012; Wang et al. 2021; Schartz et al. 2024). This increase in genomic content has been connected with an increase in nucleus size, cell size, and changes at the tissue and organ levels (Gregory 2001; Marguerat and Bähler 2012; Itgen et al. 2022). However, less research has focused on how this genomic gigantism alters the transcriptional process of the cell (Sessions and Wake 2020; Taylor et al. 2024)

In a typical eukaryotic cell, DNA is transcribed into a pre-mRNA transcript, introns are spliced out leaving only the exonic sequences, and a poly-adenylated tail is attached to make a mature mRNA transcript (Hocine et al. 2010). In the canonical form of splicing, introns are removed by spliceosomes, facilitated through small nuclear ribonucleoproteins (snRNPs) (Berget et al. 1977; Darnell 1978). The 5’ snRNP attaches to the complementary sequence of the transcript downstream of the intron; this forms a lariat structure that leads to the activation of the spliceosome, which then removes the intron all at once (Konarska et al. 1985; Sharp 1985; Gehring and Roignant 2021).

An alternative to this canonical splicing mechanism was later discovered (Hatton et al. 1998; Burnette et al. 2005). This novel form of splicing – called recursive splicing (RS) – works by removing introns in multiple sequential pieces instead of all at once (Hatton et al. 1998; Burnette et al. 2005). The mechanism works by splicing out a section of the intron flanked by discreet splicing sites, referred to as recursive splice sites (RSS) (Duff et al. 2015; Sibley et al. 2015; Kelly et al. 2015). These subsections are still removed using lariat structures and spliceosomes (Hoppe et al. 2023). As each is removed sequentially, the remaining downstream intronic sequence is brought into contact with the upstream exon at the RSS (Sibley et al. 2015; Georgomanolis et al. 2016). These ephemeral partially spliced introns can be identified through deep sequencing of total RNA (Burnette et al. 2005; Sibley et al. 2015). Their existence results in uneven RNA-seq read depths across the entire length of the intron that produce a characteristic sawtooth pattern of depth of coverage, with the RSS defining the edges of the sawtooth. This phenomenon has been identified in several model species such as fruit flies, mice, and humans (Hatton et al. 1998; Sibley et al. 2015; Joseph et al. 2018; Moon and Zhao 2022).

The presence of recursive splicing correlates with intron length; most RSS in *Drosophila* occur in introns longer than 40kb, and most RSS in mice occur in introns longer than 51kb (Pai et al. 2018; Moon and Zhao 2022). However, all this work has examined long introns in typically sized genomes. Natural genome size diversity extends across a much greater range than is seen across mammals and *Drosophila*; salamanders and lungfishes, for example, demonstrate up to a 40-fold increase in genome size relative to humans (Gregory 2024). On average, intron length in the model salamander *Ambystoma mexicanum* (genome size = 32 Gb) is 13 times longer than in humans, and similar intronic expansion in seen in the Australian lungfish *Neoceratodus forsteri* (genome size = ∼50 Gb) (Nowoshilow et al. 2018; Meyer et al. 2021). Considering this intronic expansion in species with large genomes, and the fact that RS occurs in the longest introns in previous studies of smaller genomes, we hypothesized that there would be ubiquitous reliance on recursive splicing to manage the job of removing introns in gigantic genomes.

To test this hypothesis, we looked for evidence of recursive splicing in a species with a gigantic genome. Over the last decade, advances in sequencing and assembly have made gigantic genomes amenable to genomic and transcriptomic analyses (Warren et al. 2015; Stevens et al. 2016; Nowoshilow et al. 2018; Wang et al. 2021; Schartl et al. 2024). Based on the existence of a well-annotated genome assembly and deep sequencing total RNA datasets, we selected the west African lungfish *Protopterus annectens* (40 Gb) as our model taxon. We tested some of the introns found among the 200 longest genes of *P. annectens* for evidence of recursive splicing, and we compared our findings to recursive splicing levels found among the 200 longest genes in the human genome using the same approach. Our results suggest that the largest vertebrate genomes do not rely on ubiquitous recursive splicing for removal of their spectacularly long introns.

## Materials and Methods

### Dataset Selection

We used the *P. annectens* genome assembly from Wang et al. (2021) (NCBI reference number GCA_019279795.1). In addition, we used the largest dataset for the species based on rRNA depleted RNA-seq libraries that did not use polyadenylation tail selection, as poly-A selection only selects for mature transcripts; the libraries were constructed from liver, gill, gut, and lung tissues and sequenced on an Illumina HiSeq 4000 (SRA entries SRX3392415, SRX3392416, SRX3392417, SRX3392418; ∼25 billion bp total) (Chen et al. 2021). We used the human genome as both a positive control for validating our RSS identification pipeline, and as a point of comparison representing a typically sized vertebrate genome (Sibley et al. 2015; Wan et al. 2021; Hoppe et al. 2023). We used the genome assembly used in previous human RSS analyses (GRCh37.p13, NCBI reference GCF_000001405.25) (Sibley et al. 2015). We used rRNA depleted RNA-seq libraries that did not use poly-A selection; the libraries were constructed from liver tissue, one of the tissues represented in the lungfish dataset (SRX20105218; ∼25.9 billion bp total). We subsampled the human liver RNA-seq dataset to ∼25.9 billion bp total to match the size of the lungfish RNA-seq datasets.

### Pipeline and Validation

Our pipeline followed that used by Sibley et al. (2015) to identify recursive splice sites in introns from human RNA-seq data: (1) Visually inspect reads mapped across introns for a sawtooth pattern in read depth (Sibley et al. 2015; Georgomanolis et al. 2016; Joseph et al. 2018), (2) Mark putative recursive splice site motifs in each intron (AG/GTAAG, AG/GTGAG, AG/GTAGG, AG/GTATG, AG/GTAAA, AG/GTAAT, AG/GTGGG, AG/GTAAC, AG/GTCAG, or AG/GTACG) (Sibley et al. 2015), (3) Assess whether motifs align with putative sawtooth boundaries (i.e. the center of the estimated sawtooth boundary is within 550 bp of a splice site motif; we chose this cutoff given Illumina HiSeq 4000 paired-end read lengths and the noise inherent in calculation of per-bp read depths from Illumina data), (4) Test whether segmented regression incorporating the putative recursive split site yields a significantly better-fitting relationship between intronic position and read depth than a single linear regression across the entire length of the intron. Based on our analyses following Sibley et al. (2015), we considered a ≥ 1.1 increase in slope R-squared value to support the presence of recursive splicing. We mapped RNA-seq reads to the genomic sequence using BWA and we visualized reads mapped to introns in R Studio (Allaire 2011; R Core Team 2021) using R packages dplyr (Wickman et al. 2021) and ggplot2 (Valero-Mora 2010). We used R packages parameters and effectsize to perform regression analyses (Ben-Shachar et al. 2020; Lüdecke et al. 2020).

Using this pipeline, our first goals were to (1) confirm that we could identify known RSS from human datasets, and (2) assess the depth of RNA-seq coverage necessary to identify known RSS sites. To this end, we chose three introns with strong evidence for RSS in human brain tissue (Sibley et al. 2015) that were also expressed in our human liver RNA-seq dataset: PDE4D intron #14, MAGI1 intron #22, and CADM1 intron #9. Our pipeline identified the RSS annotated in Sibley et al. 2015, confirming our pipeline’s efficacy and suggesting that these RSS sites are conserved across at least brain tissue (used in Sibley et al. 2015) and liver tissue (used here). Next, we randomly subsampled the three genes to achieve lower read depths by parsing the fastq files into smaller and smaller subsets, and we reran our pipeline on the reduced datasets. Based on the results of these subsampled datasets, we established a minimum level of sequencing coverage necessary to detect RSS: 40% of sites with > 0 reads, with no large gaps, and an average log_2_ read depth of ≥ 2.1 for the non-zero sites (Fig. 1).

**Figure 1:**
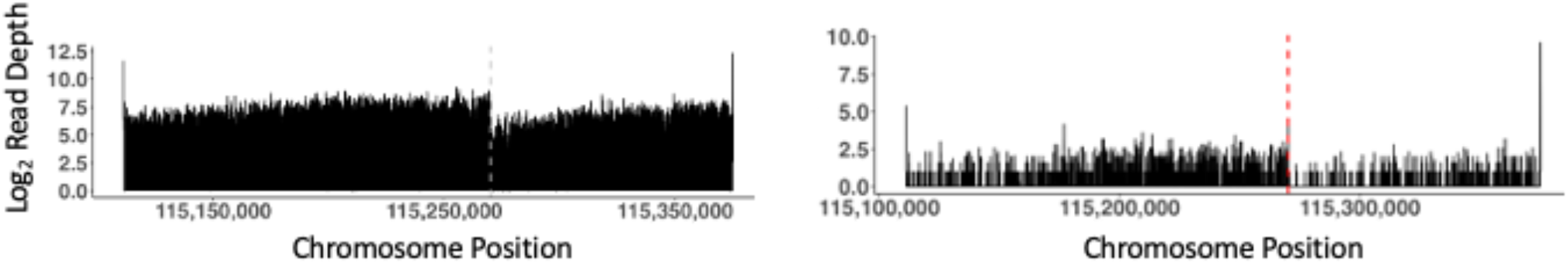
Example of RSS identification in the human CADM1 intron as pipeline validation. Left: Full RNAseq dataset. Right: Dataset subsampled to a smaller size that still allows RSS detection. The edge of the characteristic “sawtooth” in both panels is indicated with a dotted line.

**Figure 2:**
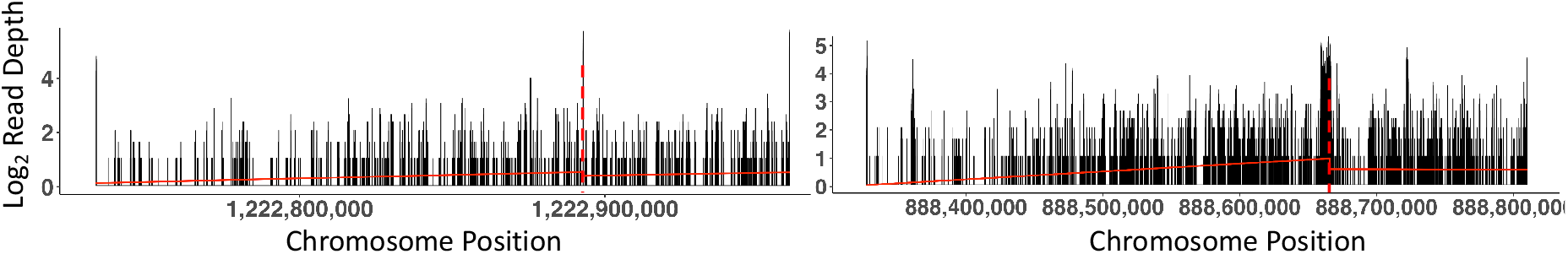
Lungfish ATAD2B Intron 15 (left) and NF1 Intron 35 (right) show RNA-seq read depth patterns consistent with removal by recursive splicing, as well as coincident RSS motif sites and improved model fit using segmented regression. Putative recursive splice sites are indicated with dotted lines.

### Testing for Recursive Splicing in Lungfish

We ordered all annotated genes in the lungfish genome from longest to shortest and selected the 200 longest genes, which ranged from 4,097,758 – 17,083,450 bp in length. Next, we obtained all of the RNA-seq dataset fastq files using the SRA Toolkit from NCBI (SRA Toolkit Development Team 2019). Twenty-five of these 200 genes had even, consistent RNA-seq coverage and were analyzed further: ARFGEF1, ATAD2B, ATP9B, BIRC6, CLIP1, CREBBP, DNMT3A, DOCK2, FAF1, FBN1, FOXP1, MTOR, NF1, PCCA, PDS5A, PPP6R2, PRKCZ, PTPRM, RANB17, REV1, RPTOR, SHANK3, TBC1D22A, UBR3, WDFY4. These 25 genes were subdivided into their 856 separate introns, which ranged in size from 85 – 1,181,814 bp in length. Of these 856 introns, 109 introns from 15 total genes passed our filter for sufficient depth of coverage to detect RSS and were moved forward through our pipeline. We also identified the 200 longest genes in the human genome and obtained recursive splicing levels for the subset of these 200 analyzed in Sibley et al. (2015) as a point of comparison.

## Results

### Recursive Splicing in Lungfish

Of the 109 introns that had sufficient depth of coverage, none showed as clear a sawtooth pattern as our subsampled human controls. However, seven showed a spike or peak in read depth within the intron that we investigated further, as we reasoned that a sawtooth edge could appear as a spike or peak in lower coverage datasets.

Of the seven introns with a spike in read depth, five introns across five genes contained an RSS motif that coincided with a putative sawtooth boundary. The other two introns contained motifs that did not coincide with any putative sawtooth boundaries. Of these remaining five introns, three showed a ≥ 1.1 increase of fit in their R-squared values under the segmented regression model with the segment defined at the putative RSS site compared to the single regression model. Further analysis of the peak sequence defining the edge of the putative sawtooth was done to see if the pattern could have been an artifact caused by a repetitive genomic sequence (e.g. a transposable element) that would have high corresponding RNA-seq depth independent of recursive splicing. One of the three sequences was identified as overrepresented in the genome, mapping to > 1000 different unplaced genome scaffolds, and was eliminated.

Overall, we found evidence consistent with recursive splicing in 1.8% (2/109) of the lungfish introns for which we had sufficient coverage to continue analysis. These two introns were from two different genes; thus, 13% (2/15) of the longest lungfish genes we analyzed showed evidence consistent with recursive splicing (Table 2).

**Table 1:** Gene and intron characteristics and regression model fits for human introns used in pipeline validation.

| Gene | Gene Length | Intron # | Intron Length | R-Squared Segmented | R-Squared Whole | Fold Difference |
| --- | --- | --- | --- | --- | --- | --- |
| PDE4D | 1,527,249 | 14 | 1,019,451 | 0.205 | 0.039 | 5.3 |
| MAGI1 | 685,393 | 22 | 684,471 | 0.485 | 0.453 | 1.1 |
| CADM1 | 335,178 | 9 | 335,178 | 0.176 | 1.19E-04 | 1,474 |

**Table 2:** Gene and intron characteristics and regression model fits for lungfish introns showing evidence consistent with removal by recursive splicing.

| Gene | Gene Length | Intron # | Intron Length | R-Squared Segmented | R-Squared Whole | Fold Difference |
| --- | --- | --- | --- | --- | --- | --- |
| ATAD2B | 4,508,040 | 15 | 227,086 | 0.023 | 0.018 | 1.28 |
| NF1 | 5,318,956 | 35 | 482,874 | 0.06 | 0.026 | 2.27 |

In the human genome, of the 200 longest genes, Sibley et al. (2015) explored 170 of them for the presence of recursive splicing, and 10% (17/170) showed evidence of an intron that undergoes recursive splicing (Sibley et al. 2015). These 170 genes contain roughly 3,794 introns – depending on which annotation is selected – thus, 0.45% (17/3,794) of those introns show evidence of recursive splicing. No genes showed evidence for more than one recursively spliced intron.

## Discussion

Sibley et al. (2015) results suggest that 0.45% of introns in the 200 longest genes in humans are removed by recursive splicing (Sibley et al. 2015). Using a similar approach, our results suggest that 1.8% of the introns in the 200 longest genes in lungfish are removed by recursive splicing. In humans, the introns that are removed by recursive splicing are always the longest intron in the gene, typically the first intron (Bradnam and Korf 2008). In contrast, in the lungfish, the introns that show evidence for removal by recursive splicing are not the longest intron in the gene, and the longest intron is not typically the first intron, suggesting a possible different relationship between recursive splicing and intron identity.

In *Drosophila*, several lines of evidence suggest that recursive splicing is functionally important for intron splicing. RSS are distributed non-randomly across the longest introns, subdividing the longest ones into equal subsections (Joseph et al. 2018; Pai et al. 2018).

Additionally, splicing becomes increasingly noisy and error-prone as intron lengths increase, and recursive splicing leads to more accurate, albeit slower, splicing of the longest introns (Pai et al. 2018). In humans, splicing also becomes noisier and more error-prone as intron length increases (Pickrell et al. 2010). However, initial estimates of recursive splicing in humans and mice – which targeted nervous system tissue, as it is characterized by longer transcript lengths – revealed lower levels than those seen in flies, despite overall longer intron lengths (Sibley et al. 2015; Joseph et al. 2018; Moon and Zhao 2022). Thus, the functional significance of recursive splicing remained obscure.

As analytical methods developed, higher levels of recursive splicing were identified in humans, revealing the phenomenon in introns of all lengths and multiple cell types beyond the nervous system (Wan et al. 2021; Hoppe et al. 2023). Importantly, live imaging of single-cell transcriptional and splicing dynamics revealed that recursive splicing is widespread, occurring in ∼30% of human genes; however, RSS selection can be stochastic, removing introns in a variety of different segments defined not at conserved sites, but rather by random selection by the spliceosome of one of many possible RSS (Wan et al. 2021). This stochastic process produces diverse, transient intermediate RNA molecules that do not appear as a sawtooth pattern in RNA-seq datasets. Taken together, this work revealed that many human genes experience recursive splicing, that it is more prevalent in longer introns, and that it can be stochastic (particularly in shorter genes) or occur constitutively at conserved RSS (Hoppe et al. 2023).

Based on these earlier studies, we hypothesized that the long introns in the gigantic lungfish genome would experience noisy, error-prone splicing if excised as single lariats, and that ubiquitous reliance on recursive splicing might have evolved to curb this noise. Our results do not support this hypothesis. In contrast, we do not see evidence for high levels of recursive splicing occurring at conserved RSS in the lungfish; 1.8% of analyzed introns show a pattern of RNA-seq read depths that bears resemblance to the predicted sawtooth as well as coincidence with an RSS motif and improved fit using segmented regression models. Given the size of our dataset and the limited number of introns for which we had sufficient sequencing depth to detect RSS, we do not place high confidence in the exact percentage of 1.8%. Increased RNA-seq read depths will certainly yield a more accurate estimate of recursive splicing levels. However, the relevant comparison is between our lungfish results and the results obtained analyzing the 13-fold smaller human genome using the same approach, which showed <1% recursively spliced introns (Sibley et al. 2015). Given the similar results for the analyses of these two genomes using a similar approach, we conclude that our results do not support the existence of ubiquitous recursive splicing in the gigantic lungfish genome. Lariat sequencing and imaging analyses will likely reveal some additional recursively spliced lungfish introns, as were revealed in humans by these technologies (Hoppe et al. 2023). We note that the lungfish sample was derived from mixed tissues, so we cannot formally exclude the possibility that high levels of recursive splicing are occurring, but at conserved sites that differ across tissue types, thus obscuring the pattern in our mixed sample. However, we consider this to be unlikely.

We emphasize that our results do not preclude the possibility of high levels of stochastic recursive splicing occurring in lungfish. In all 25 lungfish genes with sufficient coverage, we observed a pattern of decreasing RNA-seq read-depth across the length of the entire intron. Although this pattern is predicted under canonical cotranscriptional splicing, we note that it could also be produced by stochastic recursive splicing using a large number of potential RSS sites. Additional approaches to test for recursive splicing such as intron lariat sequencing and single-molecule imaging would provide additional insight into stochastic splicing (Wan et al. 2021; Hoppe et al. 2023).

In summary, although the longest introns within a species’ genome may be more likely to undergo recursive splicing, we do not find support for the hypothesis that species with extremely long introns overall show ubiquitous reliance on recursive splicing for their removal. This suggests that, even when genomes evolve to be huge, canonical splicing is able to excise the long introns with error levels that can be accommodated by the cell. Alternatively, it leaves open the possibility that stochastic recursive splicing does remove these long introns in pieces, but that the pieces are defined randomly from a large variety of potential RS sites. We advocate for additional research to reveal in more detail the ways in which genomic gigantism affects transcriptional and splicing dynamics of RNA.

## Acknowledgments

We thank members of A Adams’ thesis committee K Hoke, D Sloan, and J Hansen for helpful discussion and feedback throughout the project. Funding for this project was provided by NSF grant 1911585 to R Mueller and Colorado State University.

## Notes

### Competing Interest Statement

The authors have declared no competing interest.

